# OR1D2 receptor mediates bourgeonal-induced human CatSper activation in a G-protein dependent manner

**DOI:** 10.1101/757880

**Authors:** Yi-min Cheng, Tao Luo, Zhen Peng, Hou-yang Chen, Jin Zhang, Xu-Hui Zeng

## Abstract

During fertilization, sperm are guided towards eggs by physiological chemokines, a process named sperm chemotaxis. Human sperm chemotaxis is speculated to be mediated by olfactory receptor OR1D2 in a pathway requiring calcium influx. Bourgeonal, an artificial ligand of OR1D2, can activate CatSper, the primary calcium channel in human sperm. However, whether bourgeonal-induced CatSper activation requires OR1D2 and how CatSper is activated remain unclear. Herein, we show that OR1D2 antibody can inhibit bourgeonal-induced CatSper activation and sperm chemotaxis, proving that OR1D2 mediates bourgeonal-induced CatSper activation. Furthermore, bourgeonal-evoked CatSper currents can be greatly suppressed by either GDP-β-S or antibody of Gα_s_. Interestingly, bourgeonal can transiently increase sperm cAMP level, and this effect can be abolished by OR1D2 antibody. Consistently, bourgeonal-induced CatSper activation can be inhibited by membrane adenylate cyclases inhibitor. Overall, our results indicate that bourgeonal activates CatSper via OR1D2-G protein-cAMP pathway. Although CatSper can be activated by various physiological and environmental factors, this study represents the most recent progress proving that CatSper can be indirectly activated by extracellular regulators through a G-protein-dependent intracellular signaling pathway.

## Introduction

Mammalian sperm need to be guided towards eggs by various mechanisms, including thermotaxis (1), rheotaxis (2) and chemotaxis (3), which is defined by moving up a concentration gradient of chemokines. Mammalian sperm chemotaxis was initially described 68 years ago (4). To date, some physiological chemokines, such as progesterone (5) and atrial natriuretic peptide (6), have been identified in mammalian follicular fluid and cumulus cell secretions, suggesting that chemotaxis is a short-range mechanism acting near the fertilization site to guide sperm to the egg (7). Although the signaling pathways of mammalian sperm chemotaxis are far from being defined, one candidate, the olfactory receptor OR1D2 (olfactory receptor family 1 subfamily D member 2, aliases: hOR17-4), has been proposed to play an important role in human sperm chemotaxis (8).

Although 1.4%-4% of all human genes encode olfactory receptors (ORs) (9), two thirds of them show sequence disruption, leaving about 350 functional ORs (10, 11). Surprisingly, the expression of functional ORs is not tightly restricted to olfactory sensory neurons. For example, some ORs have been identified in early stage germ cells and mature sperm of rats and dogs (12, 13), suggesting potential roles of ORs in the mammalian reproductive system. In 2003, the mRNA transcript of OR1D2 was identified in human testis. More importantly, bourgeonal, a synthetic ligand of OR1D2, was shown to increase human sperm [Ca^2+^]_i_ (cytosolic free calcium concentration) and induce sperm chemotatic movement (8). In the follow-up studies, although other OR members (OR7A5/OR4D1) have been identified in human testis, there is no evidence supporting their involvement in the regulation of human sperm chemotaxis (14). Therefore, OR1D2 receptor remains the only candidate as a mediator of human sperm chemotaxis, although the consequent signaling after OR1D2 activation remain unclear.

In sea urchin, regulation of [Ca^2+^]_i_ fluctuations is the key feature of sperm chemotaxis (15, 16). Bourgeonal-induced human sperm chemotaxis also requires calcium influx (8), suggesting that calcium channels play key roles in human sperm chemotaxis. Although various calcium channels have been proposed to be present in human sperm (17), only CatSper, the sperm specific calcium channel, has been confirmed by patch clamping (18-21). Notably, *CatSper* gene deletion in both mice and humans produces male infertility, indicating the vital role of CatSper (22-26). It has been proposed that bourgeonal activates human sperm CatSper (27), although whether OR1D2 is necessary for bourgeonal to activate CatSper remains unclear. Since ORs usually transfer extracellular signals through G-protein dependent signaling pathways (11, 28), the involvement of molecules in the G-protein pathways should be expected in bourgeonal-mediated effects, if bourgeonal indirectly activates CatSper through binding with OR1D2. However, 250 μM GDP-β-S showed no inhibitory effect on bourgeonal-induced CatSper activation (27). In addition, no significant increase of cAMP concentration in human sperm had been detected after bourgeonal incubation (27). Those results led to the proposal that bourgeonal may activate CatSper directly (27), raising the question whether OR1D2 participates in the bourgeonal-induced human sperm chemotaxis.

Because CatSper is present in species ranging from lower invertebrates to higher mammals and plays crucial role in sperm function regulation (19, 29), the activation mechanism of CatSper has drawn intensive attention. Besides intrinsic pH and voltage sensitivities (19, 20, 21, 30, 31), CatSper can be activated by a variety of physiological and environmental factors (27, 32), which had all been proposed to activate CatSper directly, leading to the idea that CatSper is a poly-modal sensor (27). Interestingly, recently progesterone has been shown to increase CatSper current by activating orphan enzyme alpha/beta hydrolase domain containing protein 2 (ABHD2) to remove an endogenous inhibitory effect of endocannabinoid 2-arachidonoylglycerol (2-AG) on CatSper (33). Nevertheless, the studies above suggest that intracellular signaling pathways are not involved in the activation of CatSper by extracellular signaling molecules.

Apart from OR1D2, some other G-protein coupled receptors (GPCRs) have been reported to exert regulatory effects on human sperm function, for instance, CCR6 (34) and G protein-coupled receptor 18 (GPR18) (35). In addition, the presence of key components in G protein-dependent pathways, such as stimulatory subunits G_olf_, Gα_s_ and mAC (membrane adenylate cyclase) I-IX, has been confirmed by multiple techniques in human sperm (36, 37), suggesting a potential role of G-protein dependent signaling in regulating human sperm function. Thus, clarification of whether OR1D2 is necessary for bourgeonal to activate CatSper is not only critical for understanding the chemotaxis mechanism in human sperm, but may also offer insight into how CatSper is activated by extracellular stimuli in general.

In this study, OR1D2 antibody was applied to investigate whether OR1D2 is required for bourgeonal to activate CatSper and induce chemotaxis in human sperm. Furthermore, the potential involvement of intracellular signaling pathway in bourgeonal-induced CatSper activation has also been explored. Our results indicate that OR1D2 mediates bourgeonal-induced CatSper activation through a G-protein dependent manner. This study demonstrates that OR1D2-G protein-cAMP pathway is involved in human sperm chemotaxis regulation, and even more importantly, it provides an example that intracellular signaling pathways must be taken into consideration when studying the mechanisms by which CatSper is activated by extracellular stimuli.

## Results

### OR1D2 is distributed along the flagellum of human sperm

In previous studies, the expression of OR1D2 mRNA transcript in human testis (8) and the location of OR1D2 in mature human sperm (38) had already been identified. In the present study, the presence of OR1D2 in sperm was further examined by immunoblotting. In both jurkat cells (positive control) and human sperm, protein bands consistent with the control size (∼40 kD) provided by the commercial company (Sigma Aldrich) could be recognized by the OR1D2 antibody (Fig. 1A). These bands were slightly larger than the expected molecular weight of OR1D2 (∼35 kD) perhaps because of the post-translational protein modification. Similar to previous report (38), OR1D2 was distributed along the whole sperm tail (Figure S1A). In negative groups, neither fluorescence staining nor protein band could be detected after replacing OR1D2 antibody with corresponding nonspecific IgG (Figure S1A and B).

**Figure 1.**
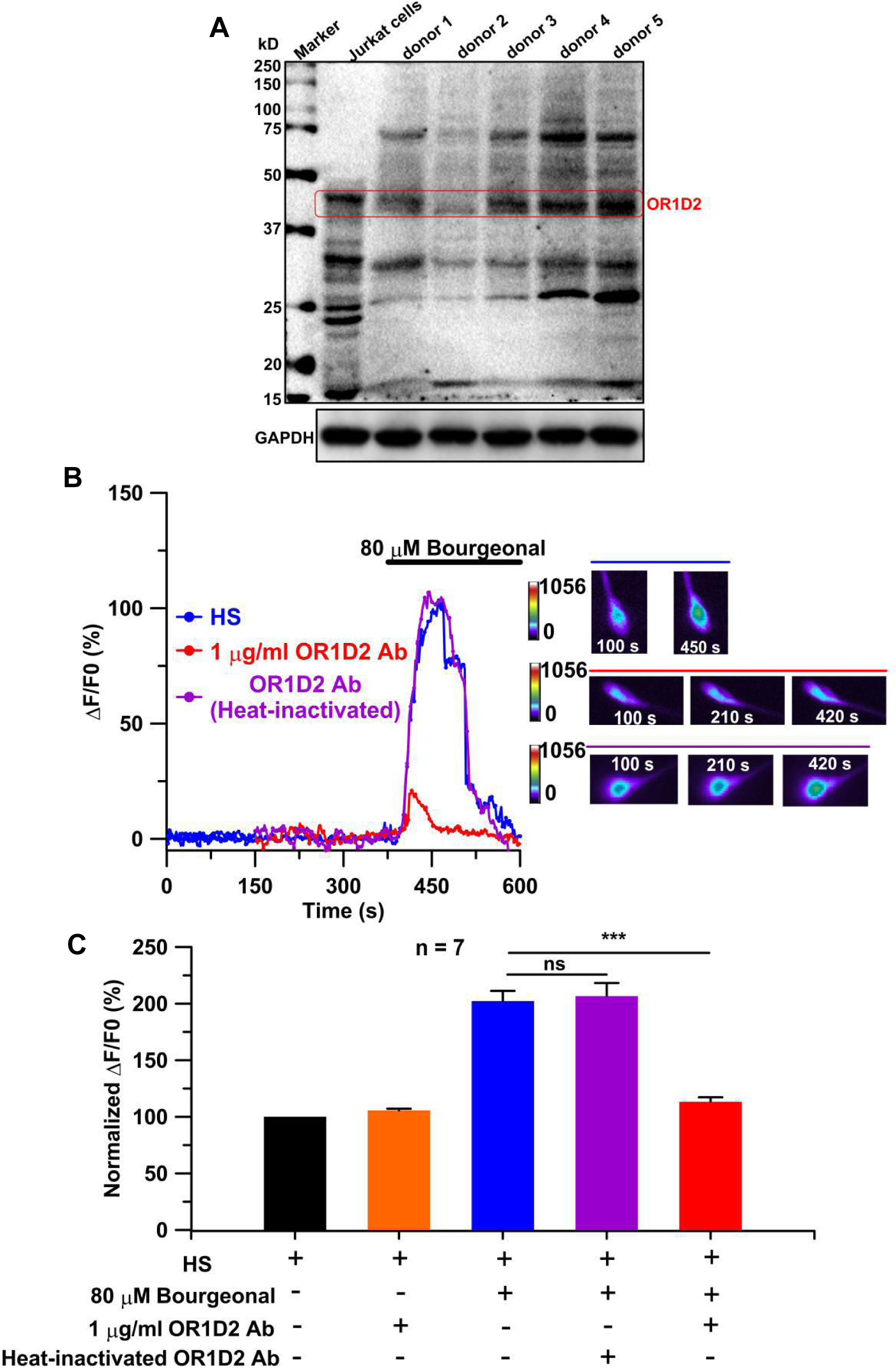
OR1D2 antibody suppresses [Ca^2+^]_i_ increase and sperm chemotaxis evoked by bourgeonal in human sperm. **(A)** The presence of OR1D2 in human sperm was validated by immunoblotting. GAPDH was used as loading control. **(B)** The effect of either untreated or heat-inactivated OR1D2 antibody on bourgeonal-induced human sperm [Ca^2+^]_i_ increase was determined by single cell calcium imaging (left panel). Examples of single cell calcium imaging were showed as in the right panel. **(C)** The influence of OR1D2 antibody on bourgeonal-evoked human sperm [Ca^2+^]_i_ increase related to (B) was analyzed. The amplitude of ΔF was calculated by subtracting resting fluorescence intensity (HS, F_0_) from maximum fluorescence intensity stimulated by various chemicals (in HS). \*\*\**P*=0.0006. Data information: Data are presented as mean ± SEM. n indicates total number of cells. Statistical differences were determined by Mann-Whitney test. \*\*\**P* < 0.001.

### OR1D2 is involved in the increase of sperm [Ca^2+^]_i_ induced by bourgeonal

To investigate whether OR1D2 is required for bourgeonal-induced CatSper activation, OR1D2 antibody was applied to examine its effect on bourgeonal-induced [Ca^2+^]_i_ increase in human sperm. Herein, single sperm calcium imaging was employed to monitor the fluctuation of sperm [Ca^2+^]_i_. Compared with bourgeonal alone, addition with OR1D2 antibody exhibited significant inhibitory effect on sperm [Ca^2^+]_i_ increase induced by bourgeonal, while application of heat-inactivated OR1D2 antibody failed to produce a similar effect (Fig. 1B and C). In addition, our results confirmed that the bourgeonal-induced [Ca^2+^]_i_ increase relies on extracellular calcium (Figure S2A) (27).

### OR1D2 mediates bourgeonal-evoked CatSper currents and chemotaxis

If OR1D2 is required for bourgeonal to activate CatSper, the interruption of OR1D2 should decrease the CatSper currents evoked by bourgeonal. To test this idea, voltage patch clamping was applied to human sperm. Consistent with the [Ca^2+^]_i_ increase after bourgeonal incubation (Figure 1B and C), bourgeonal application increased CatSper currents recorded from single human sperm (Figure 2A and B). When OR1D2 antibody was added together with bourgeonal, the increase extent of CatSper currents was significantly decreased (167.34±8.13% Vs 125.38±5.16%, Fig. 2C and D). This inhibitory effect should result from the interruption of OR1D2 by the antibody because the antibody itself did not inhibit the resting CatSper currents (Figure 2C and D). As a control, 1 μg/ml rabbit IgG had no effect on bourgeonal-induced CatSper currents (Figure S2B and C).

**Figure 2.**
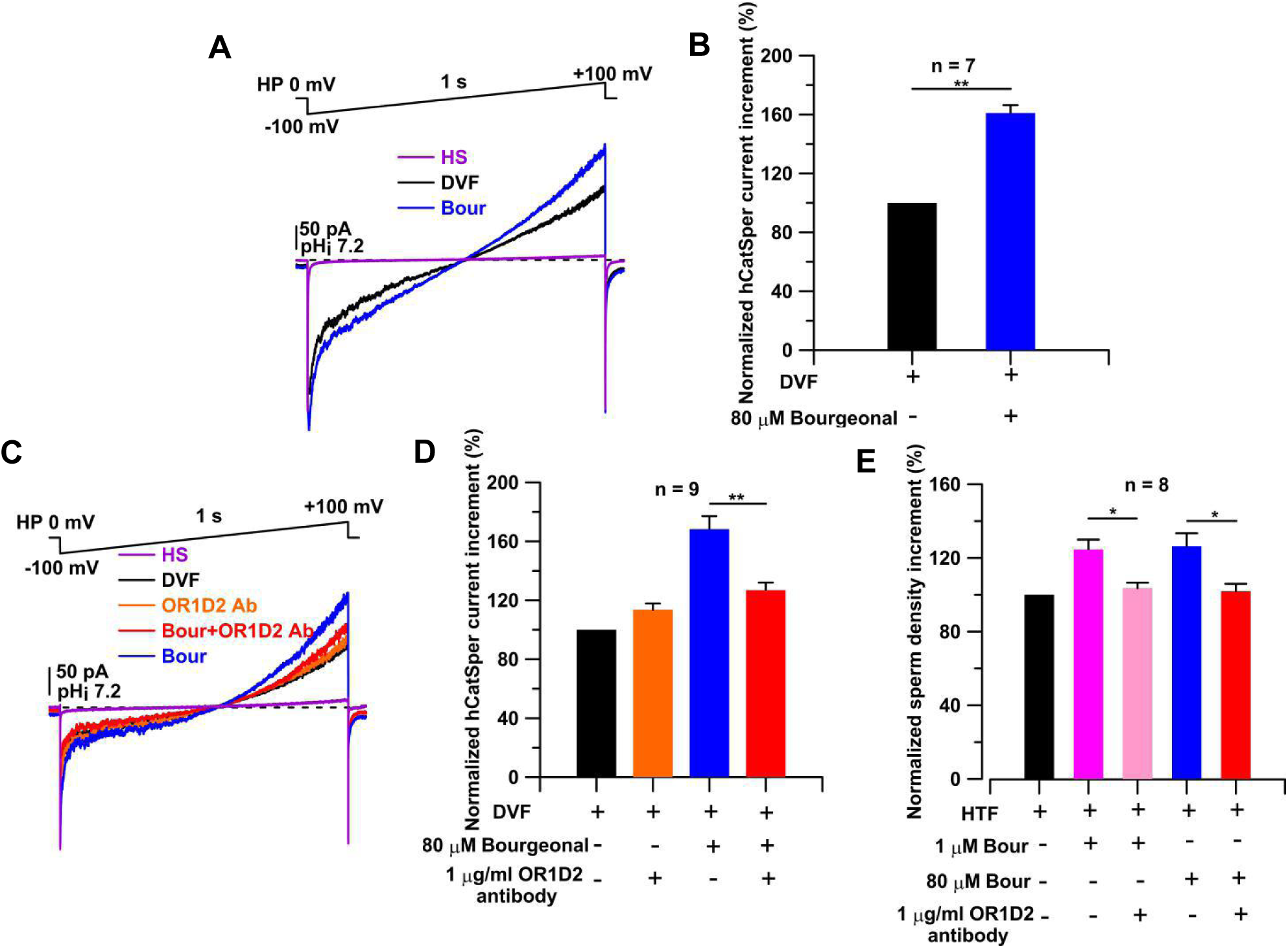
OR1D2 mediates bourgeonal-evoked CatSper currents and chemotaxis. The baselines were recorded in HS solution. Monovalent CatSper currents were recorded in divalent-free Na^+^-based bath solution (NaDVF). All chemicals were dissolved in NaDVF. **(A)** An example showed 80 μM bourgeonal induced CatSper activation by voltage-clamp ramp protocol (−100 mV to +100 mV, 1 s). The Holding Potential (HP) was 0 mV and intracellular pH was indicated by pH_i_. **(B)** The increases of CatSper currents in response to 80 μM bourgeonal at +100 mV as shown in (A) were summarized. \*\**P*=0.0014. **(C)** Bourgeonal-induced CatSper currents were compared in the presence or absence of 1 μg/ml OR1D2 antibody. **(D)** The increases in CatSper currents measured at +100 mV were summarized based on (C). \*\**P*=0.0012. **(E)** The increments of sperm density in capillaries filled with bourgeonal alone (in HTF) or the combination of bourgeonal and OR1D2 antibody (in HTF) were compared after normalizing to values from parallel, untreated controls (HTF). \**P* (left)=0.028, \**P* (right)=0.021. Data information: Data are presented as mean ± SEM. n indicates total number of independent experiments. Statistical differences were determined by Mann-Whitney test. \**P* < 0.05, \*\**P* < 0.01.

Because the specificity of OR1D2 antibody is crucial to establish the requirement of OR1D2 for bourgeonal-induced CatSper activation, the specificity of this antibody was further examined by distinct methods. Firstly, the inhibitory effect of OR1D2 antibody on bourgeonal-evoked CatSper currents was removed after inactivating the antibody by boiling for 30 minutes (Figure S3A-C). In addition, OR1D2 antibody failed to inhibit progesterone-evoked CatSper currents (Figure S4A and B), suggesting that bourgeonal and progesterone activate CatSper through distinct mechanisms. Consistent with this idea, bourgeonal-evoked CatSper currents were not be affected by either MAFP (methyl arachidonyl fluorophosphanate) or the antibody of ABHD2 (Figure S5A-D), both of which would suppress CatSper currents activated by progesterone (33). Another method to examine the specificity of OR1D2 antibody is to express OR1D2 in a heterologous system such as HEK293 cells and then validate whether this antibody inhibits the bourgeonal-evoked [Ca^2+^]_i_ increase, as previously reported (8). Surprisingly, in our hands, 500 μM bourgeonal, the same concentration as applied on the transfected HEK293 cells in the previous report, increased [Ca^2+^]_i_ in non-transfected HEK293 cells (Figure S6A and B). Therefore, the HEK293 cell system seems not suitable to further confirm the specificity of OR1D2 antibody. Nevertheless, all above favorable specificity tests support that the inhibitory effect of OR1D2 antibody on bourgeonal-induced CatSper activation should be result from the interruption of OR1D2 by the antibody. Thus, our results indicate that OR1D2 is required for bourgeonal to activate CatSper.

To further clarify the role of OR1D2 in mediating CatSper activation induced by bourgeonal, the correlation between the expressing levels of OR1D2 and bourgeonal-induced [Ca^2+^]_i_ increases was assessed. Indeed, bourgeonal caused less [Ca^2+^]_i_ increase in samples with lower OR1D2 level (Figure S7A). Furthermore, linear regression analysis from 32 donors showed that OR1D2 levels correlated positively with sperm [Ca^2+^]_i_ increases caused by bourgeonal (Figure S7B, *r*^2^ = 0.83, *P* < 0.0001). Since human sample with complete deletion of OR1D2 had not been successfully screened, the expression of OR1D2 in mouse sperm was examined. Western blot analysis showed that OR1D2 was not present in mouse sperm (Figure S8A). Accordingly, bourgeonal failed to increase mouse CatSper currents, which could be activated by the well-established stimulus sodium bicarbonate (NaHCO3) (19, 39) (Figure S8B and C). Overall, these additional evidences confirm that OR1D2 mediates bourgeonal-evoked CatSper activation.

In fact, the idea that OR1D2 receptor mediates human sperm chemotaxis was solely based on the previous report that the OR1D2 agonist bourgeonal could evoke sperm chemotaxis (8). To further confirm the role of OR1D2 in mediating human sperm chemotaxis, the effect of OR1D2 antibody on bourgeonal-evoked sperm chemotaxis was examined. Human sperm chemotaxis was assessed by capillary tube method. Because higher bourgeonal concentrations were usually used to induce CatSper activation (27) while lower concentrations were used to examine chemotaxis (8), we examined both lower and higher concentrations of bourgeonal. In our hands, both 1 and 80 μM bourgeonal stimulated a similar extent of sperm accumulation in capillaries, and the effect of either concentration of bourgeonal could be abolished by 1 μg/ml OR1D2 antibody (Fig. 2E), confirming the role of OR1D2 in mediating human sperm chemotaxis.

### G proteins are involved in bourgeonal-induced CatSper activation

Since OR1D2 belongs to the family of classic G-protein coupled receptors (8, 11), the involvement of G proteins in OR1D2 mediated CatSper activation by bourgeonal was examined. As the first test, 3 mM GTP was added in the pipette solution to evaluate its effect on bourgeonal-induced CatSper currents. Although the addition of GTP did not change the basal I_*CatSper*_ densities and the final extent of increase caused by bourgeonal (Figure S9A and B), it increased the slope parameter of CatSper activation process (Figure 3A and B), suggesting the involvement of G proteins in bourgeonal-induced CatSper activation. Furthermore, GDP-β-S (Guanosine 5’-[β-thiol] diphophate), which is usually used to block G-protein dependent signaling pathway (40-42), was employed in the pipette solution to examine the involvement of G proteins. 250 μM GDP-β-S showed no effect on the increase of *I*_*CatSper*_ densities induced by bourgeonal (Figure S9C and D), consistent with previous report (27). However, the sensitivity of CatSper to bourgeonal was almost abolished by 1 mM GDP-β-S (Fig. 3C and D, Figure S9E), which could completely suppress GTP-induced G protein transducin (G_t_) activation in vertebrate retina rods and cones (43). Worth of note, the addition of 1 mM GDP-β-S in pipette solution also did not change the basal *I*_*CatSper*_ densities (Figure S9E). To examine the specificity of 1 mM GDP-β-S on G protein-dependent signaling pathways, the effect of 1 mM GDP-β-S on CatSper activation caused by 10 mM NH4Cl was validated and no inhibitory effect was observed (Figure S10A and B). Because NH4Cl activates CatSper channels via alkalizing intracellular pH (20, 21), a mechanism independent of G proteins, these results support the involvement of G-proteins in bourgeonal-induced CatSper activation.

**Figure 3.**
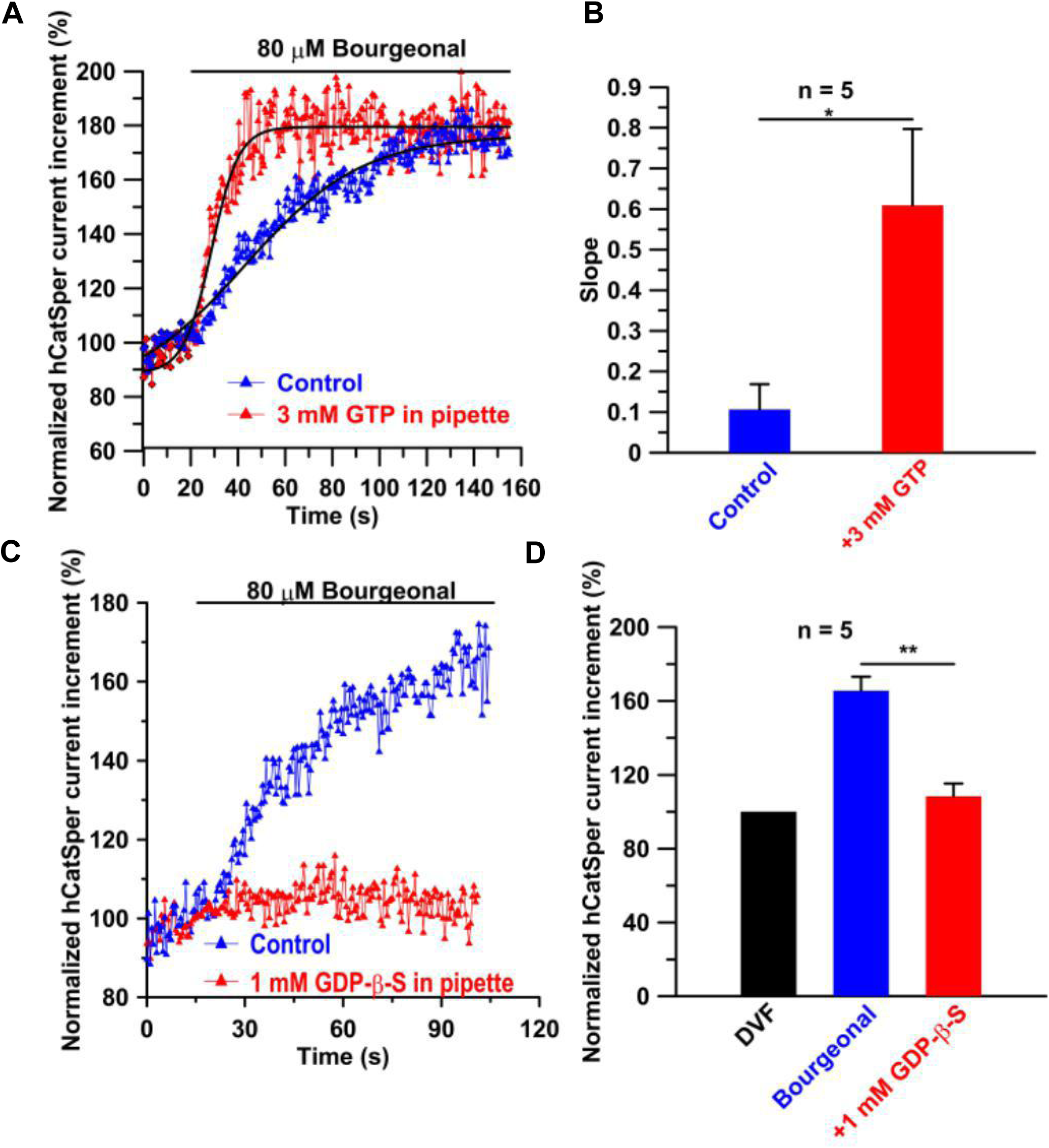
G proteins may be involved in bourgeonal-induced CatSper activation. **(A)** Bourgeonal-induced CatSper activation in the presence or absence of intracellular GTP (3 mM) at +100 mV had been compared. The data were obtained from the same donor. The activation processes were fitted by sigmoidal function, and the slope parameters were used to reflect difference in the time course of CatSper activation under both conditions. **(B)** The slope parameters of bourgeonal-induced CatSper activation in the presence or absence of intracellular GTP were summarized based on (A). *\*P*=0.016. **(C)** An example showed the effect of intracellular GDP-β-S (1 mM) on CatSper currents induced by bourgeonal. The data were recorded at +100 mV from the same donor. **(D)** The effects of GDP-β-S on bourgeonal-induced CatSper activation were summarized based on (C). \*\**P*=0.0079. Data information: In B and D, data are presented as mean ± SEM. n indicates total number of independent experiments. Statistical differences were determined by Mann-Whitney test. \**P* < 0.05, \*\**P* < 0.01.

### Stimulatory subunit Gα_s_ plays a role in bourgeonal-induced CatSper activation

To assess the potential role of different G protein signaling pathways in bourgeonal-induced CatSper activation, antibodies against different subunits of G proteins were added in the pipette solution while recording CatSper currents. Because OR1D2 is an olfactory receptor and the expression of G_olf_ subunit in human sperm had been detected (36, 37), the effect of G_olf_ antibody on bourgeonal-induced CatSper currents was evaluated. However, the addition of 2 μg/ml G_olf_ antibody showed no effect on bourgeonal-induced CatSper activation (Figure 4A and B). Because Gα_s_ had also been detected in human sperm (36, 37), we then tested Gα_s_ antibody in this type of experiment. Gα_s_ antibody greatly attenuated the CatSper currents evoked by bourgeonal (169.8±21.3% VS 115.5±9.3%, Fig. 4C and D). Furthermore, intracellular Gα_s_ antibody did not inhibit basal *I*_*CatSper*_ densities (Figure S11A), excluding the possibility that the observed attenuation was caused by direct inhibition of Gα_s_ antibody on CatSper channels. As a negative control, heat-inactivated Gα_s_ antibody failed to suppress bourgeonal-induced CatSper currents (Figure S11B). Because the primary structures of G_olf_ and Gα_s_ subunits are similar, the specificity of Gα_s_ antibody was checked by confirming that the antigenic epitope of the Gα_s_ antibody is not present in the sequence of G_olf_ subunit, precluding the possibility that Gα_s_ antibody acts on the G_olf_ subunit. Since G proteins are composed of Gα and Gβγ subunits (44), Gue1654 (45), a selective inhibitor of Gβγ, was applied intracellularly to examine whether Gβγ is involved in bourgeonal-induced CatSper activation, and the results suggested that Gβγ is not important in that process (Figure S11C-E). Overall, these results support the idea that Gα_s_ is a key component of the signaling pathway that mediates the effect of bourgeonal on CatSper.

**Figure 4.**
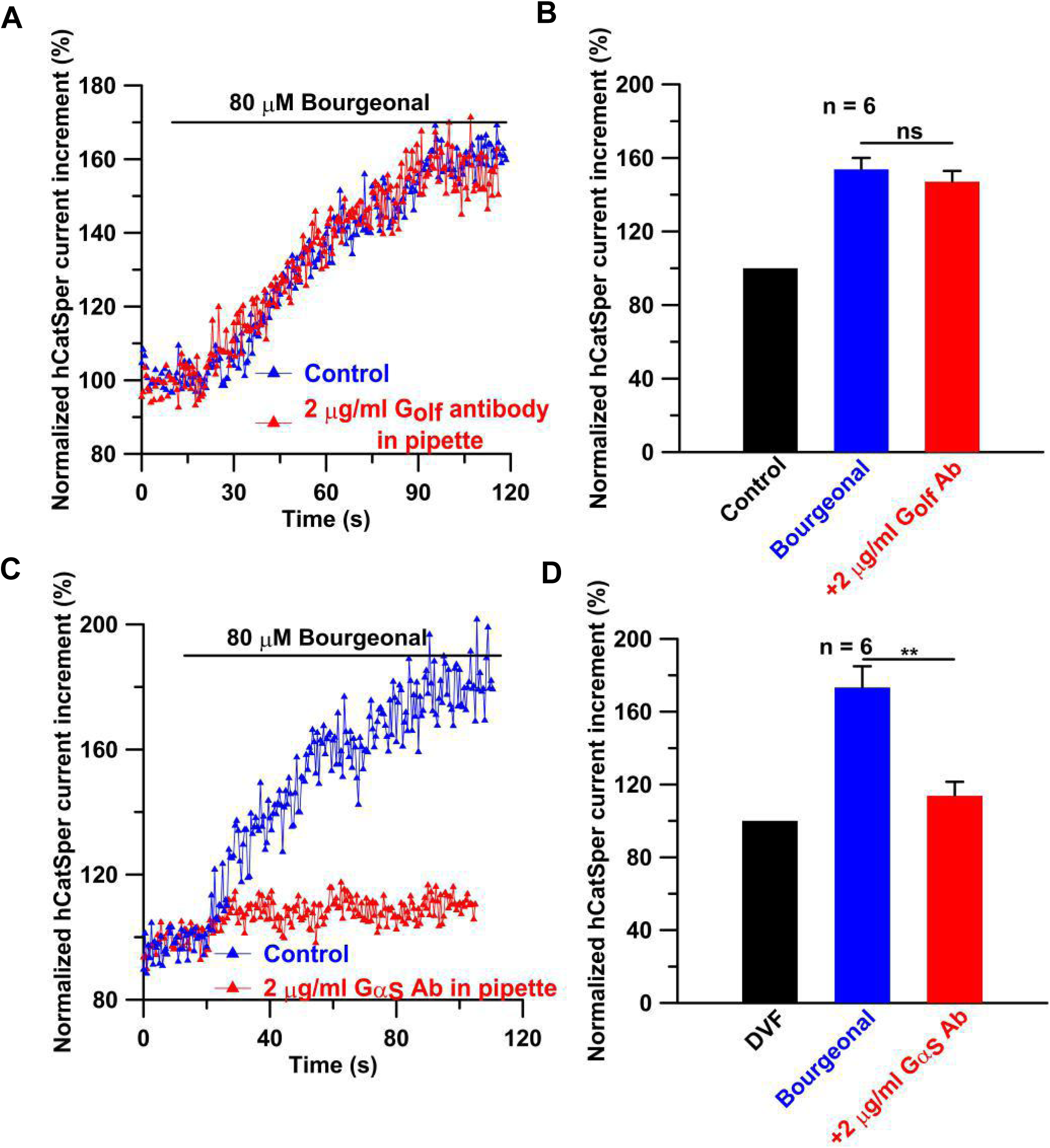
The antibody of Gα_s_ rather than G_olf_ inhibits bourgeonal-induced CatSper activation. **(A)** Contrast examples of bourgeonal-evoked CatSper currents with or without intracellular G_olf_ antibody (2 μg/ml) were showed. **(B)** G_olf_ antibody showed no significant effect on bourgeonal-evoked CatSper currents based on (A). **(C)** Examples of bourgeonal-evoked CatSper currents with or without intracellular Gα_s_ antibody (2 μg/ml) were exhibited. The currents were obtained from the same donor for comparison. **(D)** The statistical effect of intracellular Gα_s_ antibody on bourgeonal-induced CatSper currents based on (C) was examined. \*\**P*=0.0087. All currents were recorded at +100 mV. Data information: Data are presented as mean ± SEM. n indicates total number of independent experiments. Statistical differences were determined by Mann-Whitney test. ns = not significant, \*\**P* < 0.01.

### Bourgeonal transiently increases cAMP levels in human sperm

If Gα_s_ plays a key role during the process of CatSper activation caused by bourgeonal, an increase of cAMP concentration is expected in that process. However, in the previous report (27), no significant increase was detected. To re-examine this issue, Cayman’s cAMP assay, a very sensitive competitive ELISA that permits cAMP measurements within a wider concentration range (0.3-750 pmol/ml, compared to the range of 1.5-450 pmol/ml in previous report (27)), was utilized in this study. To evaluate the utility of this method, we first examined the ability of NaHCO3, a known stimulant of the soluble adenylate cyclase (sAC) activity in sperm, to increase cAMP in human sperm samples. A significant increase of cAMP levels could be observed after at least 20 s of sodium bicarbonate incubation (Figure 5A), confirming the reliability of this method. When incubating human sperm with bourgeonal for different time periods, an apparent increase of cAMP levels could be observed after 20 s, with the increase at 30 s exhibiting significance (increased from 53±3.8 to 76±5.1 pmol cAMP/10^8^ sperm, Figure 5B). For longer incubation periods, cAMP concentrations declined to the levels near resting state (Figure 5B). More importantly, the increase of cAMP concentrations induced by bourgeonal was greatly suppressed by OR1D2 antibody (Fig 5C). In contrast, CatSper inhibitor mibefradil, which could suppress the increase of CatSper current caused by bourgeonal (Figure S12A and B) (27), exerted no influence on the levels of cAMP enhanced by bourgeonal (Figure 5D), indicating the increase of cAMP is an event before CatSper activation. An obvious increase of cAMP concentrations caused by phosphodiesterase inhibitor isobutylmethylxanthine (IBMX) also confirmed the reliability of the method used here (Figure 5C and D). All of these results support the idea that bourgeonal produces an increase of cAMP concentration, although the increase appears transient.

**Figure 5.**
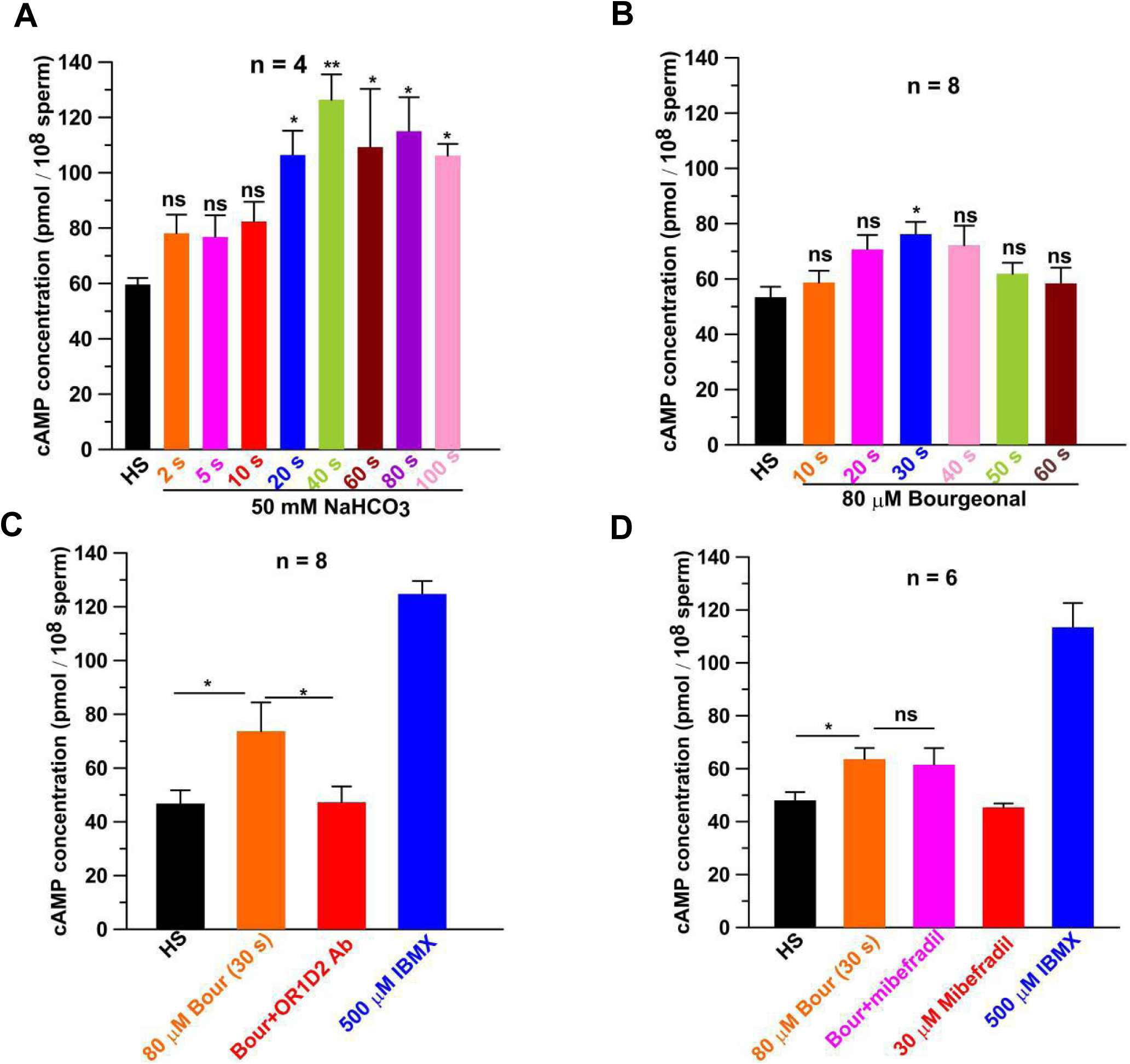
Bourgeonal transiently increases cAMP levels in human sperm. **(A)** The concentrations of sperm cAMP were examined after incubating sperm with 50 mM sodium bicarbonate (NaHCO_3_) for different periods. **(B)** cAMP concentrations in human sperm were detected after treatment with 80 μM bourgeonal for different periods. **(C)** The effect of 1 μg/ml OR1D2 antibody on bourgeonal-evoked increase of human sperm cAMP was determined. IBMX, a well known phosphodiesterase inhibitor, was used as the positive control. *P*\*(HS vs Bourgeonal)=0.039, *P*\*(Bourgeonal+OR1D2 antibody vs Bourgeonal)=0.049. **(D)** Bourgeonal-evoked cAMP increase was not affected by CatSper inhibitor mibefradil (30 μM). *P*\*(HS vs Bourgeonal)=0.015. In both C and D, the cAMP levels were detected at 30 seconds after incubation of the corresponding chemicals listed. Data information: Data are presented as mean ± SEM. n indicates total number of independent experiments. Statistical differences between untreated controls (HS) and treated groups in A and B were determined by Newman-Keuls Multiple comparison test, statistical differences of data from C and D are analyzed by students unpaired t-test. ns = not significant, \**P* < 0.05.

### mACs are responsible for the cAMP increase caused by bourgeonal

To distinguish which kind of adenylate cyclases are responsible for the cAMP increase caused by bourgeonal, inhibitors of mACs (membrane adenylate cyclases) and sAC (soluble adenylate cyclase) were utilized in this study. MDL12330A (*cis*-N-(2-phenylcyclopentyl)-azacyclotridec-1-en-2-amine hydrochloride) and SQ22536 (9-tetrahydro-2′-furyl adenine) were utilized to examine the involvement of mACs. Although 100 μM MDL12330A was able to abolish the activity of mACs (46), MDL12330A at this concentration has also been shown to inhibit CatSper channels significantly (27), a result confirmed here (Figure S13A). Because SQ22536 has been shown to inhibit the increase of sperm [Ca^2+^]_i_ caused by bourgeonal with an IC_50_ of 2 mM (37), 3 mM SQ22536 was used initially. However, SQ22536 at this concentration blocked CatSper currents significantly (Figure S13B). In addition, SQ22536 above 200 μM increased the cAMP levels in human sperm (Figure S13C) (27). Therefore, to minimize such confounding effects of SQ22536, we had utilized 100 μM SQ22536, a concentration that neither inhibited CatSper currents (Figure 6A and B), nor increased cAMP levels (Figure S13C). 100 μM SQ22536 significantly suppressed CatSper activation induced by bourgeonal (181.8±12.8% VS 106.4±4.9%, Figure 6A and B). Similarly, 100 μM SQ22536 substantially attenuated the increase of [Ca^2+^]_i_ caused by bourgeonal (Figure 6C and D). In contrast, KH7, an inhibitor of sAC, showed no effect on the increases of either CatSper currents or [Ca^2+^]_i_ caused by bourgeonal (Figure S14A-D), although the concentration used here is expected to attenuate the activity of sAC (47, 48). Together, these results indicate that mACs are responsible for the cAMP increase caused by bourgeonal.

**Figure 6.**
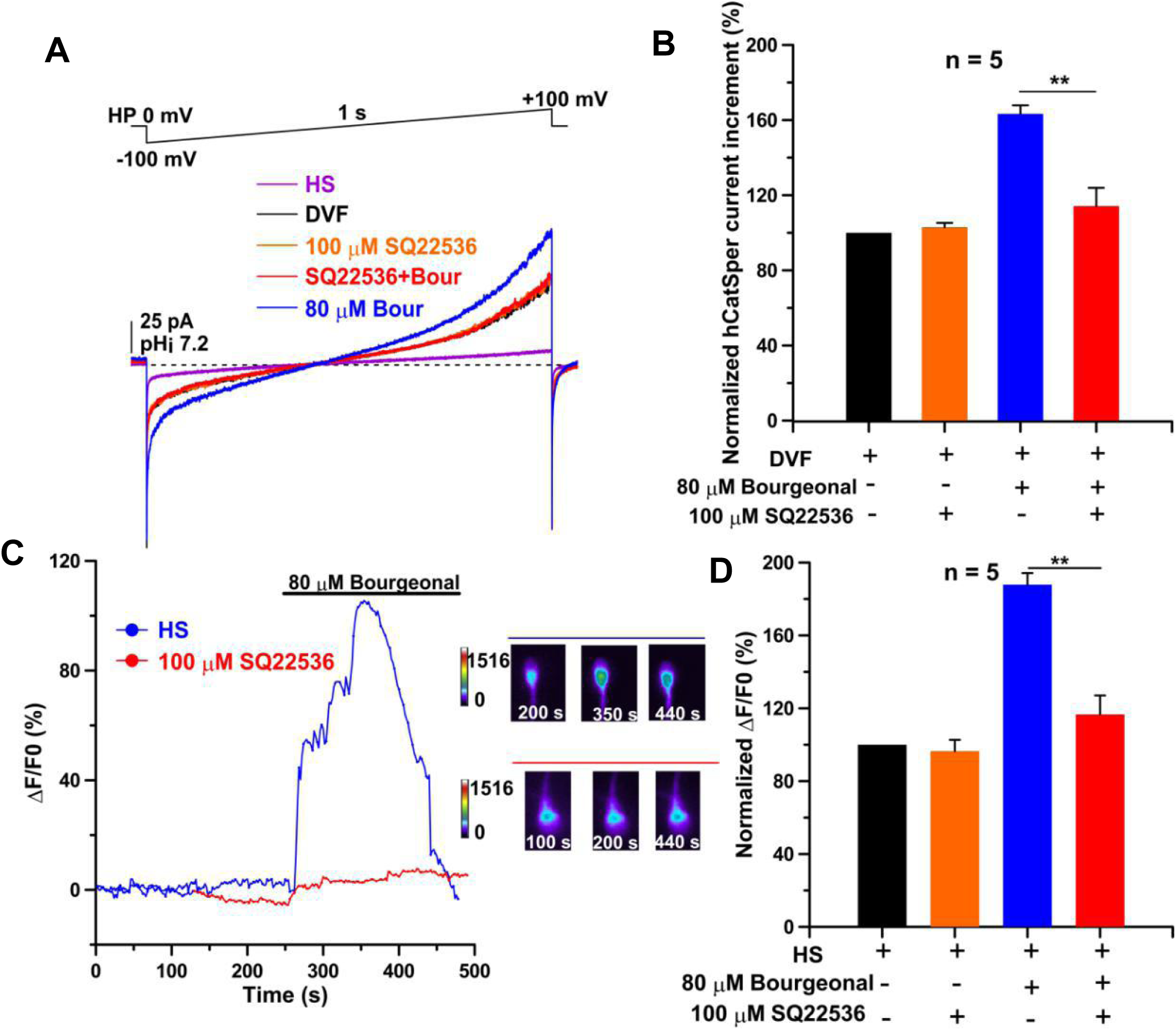
mACs inhibitor SQ22536 attenuates bourgeonal-induced human CatSper activation. **(A)** CatSper current examples were recorded in response to SQ22536 and bourgeonal. **(B)** The relative inhibition of SQ22536 on bourgeonal-evoked CatSper currents was examined based on (A). Current amplitudes at +100 mV were compared. \*\**P*=0.0079. **(C)** The inhibitory effect of SQ22536 on bourgeonal-induced [Ca^2+^]_i_ increase in human sperm was showed by calcium imaging. **(D)** The influence of SQ22536 on bourgeonal-evoked sperm [Ca^2+^]_i_ increase was analyzed based on (C). \*\**P*=0.0081. Data information: In B and D, data are presented as mean ± SEM. n indicates total number of independent experiments / total number of cells. Statistical differences were determined by Mann-Whitney test. \*\**P* < 0.01.

## Discussion

To clarify the role of OR1D2 in human sperm chemotaxis, OR1D2 antibody was utilized to investigate whether OR1D2 is required for bourgeonal to activate CatSper and induce chemotaxis in human sperm. Our results showed that OR1D2 antibody significantly inhibited the increase of [Ca^2+^]_i_, the amplification of CatSper currents, and chemotaxis induced by bourgeonal, confirming that OR1D2 is involved in bourgeonal induced CatSper activation and chemotaxis in human sperm. Furthermore, the results proved that OR1D2 mediates bourgeonal-induced CatSper activation in a G-protein dependent manner, because the effect of bourgeonal on CatSper currents could be reduced by intracellular addition of either GDP-β-S or Gα_s_ antibody. In addition, the increase of cAMP could be detected after bourgeonal application, and this increase could be abolished by OR1D2 antibody. Consistently, SQ22536, an inhibitor of mACs, suppressed CatSper activation induced by bourgeonal. Thus, this study not only establishes a relatively intact signaling pathway to explain human sperm chemotaxis but also provides the first example showing that CatSper can be activated by extracellular stimulus through an intracellular signaling pathway.

### OR1D2 is a key component in the signaling pathway during bourgeonal-induced human sperm chemotaxis

Although the underlying mechanism of mammalian sperm chemotaxis remains largely unknown, human sperm chemotaxis is speculated to be mediated by the olfactory receptor OR1D2, a concept solely relying on the previous report that OR1D2’s agonist bourgeonal could evoke sperm chemotaxis (8). Bourgeonal-induced human sperm chemotaxis required the increase of [Ca^2+^]_i_ (8), and the increase of human sperm [Ca^2+^]_i_ was resulted from the activation of CatSper by bourgeonal (27). However, 250 μM GDP-β-S showed no inhibitory effect on bourgeonal-induced CatSper activation (27). In addition, no significant increase of cAMP concentration in human sperm had been detected after bourgeonal incubation (27). Those results supported the claim that bourgeonal may activate CatSper directly (27). In the other word, bourgeonal may induce human sperm chemotaxis without the involvement of OR1D2, raising the question whether OR1D2 is a key mediator required for human sperm chemotaxis.

After confirming the expression of OR1D2 in human sperm by western blot analysis (Figure 1A), this study found that OR1D2 antibody showed inhibitory effect on bourgeonal-induced sperm [Ca^2+^]_i_ increase (Figure 1B and C), the amplification of CatSper currents (Figure 2C and D), and sperm chemotactic movement (Figure 2E). These results support that OR1D2 is required for bourgeonal-induced chemotaxis in human sperm.

Except for antibody application, currently there is no specific method to interrupt OR1D2 in human sperm. During this study, we were fully aware that the conclusions are highly dependent on the specificity of the OR1D2 antibody. Accordingly, the specificity of the inhibition of OR1D2 antibody on bourgeonal-caused CatSper activation was examined by multiple approaches. First, the inhibitory effect of OR1D2 antibody on bourgeonal-evoked CatSper currents was ablated after denaturing the antibody by boiling (Figure S3A-C). Furthermore, OR1D2 antibody failed to inhibit progesterone-evoked CatSper currents (Figure S4A and B), while bourgeonal-evoked CatSper currents were not be affected by either MAFP or the antibody of ABHD2 (Figure S5A-D), both of which would suppress CatSper currents activated by progesterone (33), strongly arguing that OR1D2 antibody specifically interrupt CatSper activation caused by bourgeonal. Finally, the expression levels of OR1D2 and [Ca^2+^]_i_ increases caused by bourgeonal exhibited significant positive correlation (Figure S7B, *r*^2^ = 0.83, *P* < 0.0001). Overall, all of these results support that the antibody utilized here would specifically interrupt OR1D2.

During this study, OR1D2 antibody was applied from the extracellular side, thus it is important to consider the epitope of the antibody and the topology of OR1D2. According to the information provided by Sigma Aldrich, the epitope of this OR1D2 antibody covers amino acids 201-250. Based on transmembrane prediction analysis by TMpred (https://embnet.vital-it.ch/software/TMPRED_form.html), these amino acids span the 5-6^th^ transmembrane regions, and the amino acid sequence (aa223-236) connecting the 5-6^th^ transmembrane domains may locate either outside or inside of the membrane. Although UniProt database adopts the prediction that aa223-236 are located in the cytoplasm perhaps because this inside-model scores slightly higher than the outside-model (12976 vs 11068), the interruption effect of the antibody on OR1D2 showed in this study support that this 5-6^th^ linker is located outside of the membrane. Consistent with our prediction, the topology of Drosophila olfactory receptors may vary from the classic GPCRs by locating their N-terminus intracellularly rather than extracellularly and predicting the accessibility of the 5-6^th^ linker from the extracellular side (49). Anyway, although figuring out the topology of olfactory receptors is not the purpose of this study, our results strengthen the requirement of caution when comparing the topology of human olfactory receptors with classic GPCRs. Nevertheless, we also recognize the limitation of drawing conclusions mainly based on antibody interruption although currently there is no better method to interrupt OR1D2 in human sperm.

Apart from OR1D2, transcripts of other ORs mRNA could also be detected in human testicular tissue, such as OR7A5, OR4D1 and OR1D5 (aliases: hOR17-2) (8, 14). Since OR7A5 and OR4D1 were supposed not involving in human sperm chemotaxis (14), the presence and possible role of OR1D5 in mediating bourgeonal-induced CatSper activation was examined here. Immunoblotting assay suggested the existence of OR1D5 in human sperm (Figure S15A). However, OR1D5 antibody had no effect either on the increase of CatSper currents (Figure S15B and C) or on the sperm chemotaxis (Figure S15D) caused by bourgeonal. These results strengthen the importance of OR1D2 in mediating bourgeonal-induced sperm chemotaxis. However, since bourgeonal is an artificial ligand of OR1D2, the possibility that other olfactory receptors might involve in human sperm chemotaxis under physiological conditions can not be completely excluded.

### Bourgeonal/OR1D2 activates CatSper via a G-protein dependent manner

Since the activation of olfactory receptors usually involves G-protein dependent signaling pathway, the failure for 250 μM GDP-β-S to affect bourgeonal-induced CatSper activation and the failure to detect the cAMP increase after bourgeonal application are puzzling (27). In this study, a series of experiments were designed to investigate whether G-protein dependent pathway involves in CatSper activation caused by bourgeonal. First, addition of 3 mM GTP in the pipette solution could accelerate the activation process of bourgeonal on CatSper currents (Figure 3A and B). Second, although 250 μM GDP-β-S did not affect bourgeonal-induced CatSper activation (Figure S9C and D), increasing the concentration of GDP-β-S to 1 mM, which could completely suppress the activation of G_t_ caused by GTP in vertebrate retina rods and cones (43), definitely showed obvious inhibitory effect (Figure 3C and D), without affecting the basal or NH4Cl-induced CatSper currents (Figure S9E, S10A and B). Third, addition of Gα_s_ antibody in the pipette solution largely inhibited bourgeonal-induced CatSper activation (Figure 4C and D). More importantly, by utilizing a sensitive assay method, an increase of cAMP could be detected after bourgeonal application (Figure 5B), and this increase could be abolished by OR1D2 antibody (Figure 5C), but not by CatSper inhibitor mibefradil (Figue 5D). Finally, the effects of bourgeonal could be inhibited by low concentration of SQ22536 (100 μM), the inhibitor of mAC (Figure 6). Taking these together, our results strongly argue that bourgeonal/OR1D2 activates CatSper via a G-protein dependent manner.

This study reveals that OR1D2 signaling cascade depends on Gα subunit, which is weakly membrane bound, raising the possibility that Gα subunit may be removed from the cytosol upon perfusion with the pipette solution. To testify this possibility, after whole cell configuration formation, CatSper currents were recorded with repeated bourgeonal applications. In a patch successfully lasting for 45 minutes, the extent of bourgeonal-induced CatSper current amplification declined 15 minutes later (Figure S16A-D). However, the effects of bourgeonal within 5 minutes were stable (Figure S16E). In addition, amphotericin perforated-patch recording (18) was applied to compare with the regular whole-cell configuration. Within 5 minutes after whole-cell formation, these two recording modes detected similar extents of bourgeonal-induced CatSper current amplification (Figure S16F and G). The results above suggest that, although a dilution of cytosolic Gα subunit during regular whole cell mode is possible, it should not affect the effect of bourgeonal on CatSper activation within 5 minutes after whole cell formation. It is worth noting that the CatSper currents were usually recorded within 5 minutes after the formation of whole cell configuration in this study.

In contrast to the failure of detecting cAMP increase in the previous report (27), this study successfully detected a significant cAMP increase after bourgeonal application (Figure 5B). This inconsistence may be based on two facts. First, different detection methods were used in these two studies, and the method used here has wider measurement range (0.3-750 pmol/ml VS 1.5-450 pmol/ml). Second, the increase of cAMP is intrinsically transient. In fact, the increase of cAMP only exhibited significance after 30 s of bourgeonal application (Figure 5B), making it very possible to miss the “right” time point. It should be taken seriously that the increase in sperm cAMP level caused by either IBMX or NaHCO3 greatly exceeded that caused by bourgeonal, thus the effects of IBMX and NaHCO3 on human CatSper should be examined in future to further validate the linkage between cAMP and CatSper activation.

This research shows that 3 mM SQ22536 is not suitable to test whether mACs are involved in bourgeonal-induced CatSper activation because SQ22536 inhibited CatSper significantly (Figure S13B). In addition, SQ22536 above 200 μM increased the cAMP levels in human sperm (Figure S13C) (27). However, at the concentration of 100 μM, SQ22536 neither inhibited CatSper currents (Figure 6A and B) nor affected resting cAMP level (Figure S13C). Interestingly, 100 μM SQ22536 substantially abolished bourgeonal-induced CatSper activation (Figure 6A and B) and the increase of [Ca^2+^]_i_ (Figure 6C and D), indicating the activation of mACs following bourgeonal application. Since some studies pointed out that sAC rather than mACs may be the key factor for cAMP regulation in some mammalian sperm (47), the effect of KH7, the inhibitor of sAC had been examined and no effect on the increases of either CatSper currents or [Ca^2+^]_i_ caused by bourgeonal was observed (Figure S14A-D), further strengthening the role of mACs in the process of bourgeonal induced CatSper activation.

It is surprise that Gα_s_ rather than G_olf_ appears to be the key G-protein component in mediating the effect of bourgeonal on CatSper (Figure 4). Since the epitope of Gα_s_ antibody dose not present in the sequence of G_olf_ subunit, the possibility that Gα_s_ antibody acts on G_olf_ subunit should be low. Whether Gα_s_ or other unidentified Gα subunit is responsible for mediating bourgeonal-induced CatSper activation definitely requires further investigation. Nevertheless, all of the results indicate that G-protein dependent signaling pathway is responsible for CatSper activation caused by bourgeonal/OR1D2.

Because of its crucial roles in mammalian sperm function, the activation mechanism of CatSper has drawn intensive attention. Besides intrinsic pH and voltage sensitivities (19, 20, 21, 30, 31), CatSper is able to be activated by many physiological and environmental factors (27, 32). However, almost all previous studies suggested that intracellular signaling pathways are not involved in the activation of CatSper by extracellular signaling molecules. For example, CatSper may be activated as a poly-modal sensor (27), or amplified by removal of CatSper blocker (33). In contrast, this study provides an example showing that CatSper can be activated through the G-protein dependent signaling pathway. Interestingly, a most recent study found that cAMP could increase CatSper currents in mouse sperm (50). Because some other GPCRs have been reported to exert regulatory effects on human sperm functions, such as CCR6 (34) and GPR18 (35), whether G-protein dependent manner is a common signaling pathway to activate or regulate CatSper is worthy for further investigation.

### What is the downstream pathway after cAMP increase to activate CatSper channel in human sperm?

CatSper inhibitor mibefradil showed no inhibitory effect on bourgeonal-induced cAMP increase (Figure 5D), consistent with the idea that CatSper activation is a downstream event after cAMP increase. The regulatory effects of cAMP on cellular functions are usually attributed to the activation of protein kinase A (PKA) (50-52). Thus, the roles of PKA in bourgeonal-induced CatSper activation were examined in this study. Three kinds of membrane-permeable selective inhibitors of PKA (H89, KT5720 and RpcAMP) were utilized (53-56), and all of these three inhibitors could significantly inhibit bourgeonal-induced CatSper activation while not affecting basal CatSper activity (Figure S17A-H), suggesting that PKA may be involved in the activation of CatSper induced by bourgeonal. However, how PKA activation resulting in CatSper channel opening definitely requires further exploration.

Apart from PKA, cAMP can also exert its effects on cellular functions via exchange proteins directly activated by cAMP (EPACs) (57). EPACs are a family of the guanine nucleotide exchange factors, which consist of two isoforms: EPAC1 and EPAC2 (57). In general, EPACs couple cAMP production to the activation of Rap, a small molecular weight GTPase of the Ras family (58). It is worth noting that the EPACs activation can impose profound influences on ion channels, and thereby regulate a broad array of cellular physiological processes such as cell adhesion (59, 60). Interestingly, the presence of EPACs and their functional roles in mobilizing acrosomal calcium during sperm exocytosis have already been reported (61, 62). Thus, it will be necessary to determine whether EPACs are also involved in the CatSper activation and human sperm chemotaxis induced by bourgeonal.

## Materials and methods

### Chemicals

Reagents used in this study were from the following sources: bourgeonal (Enzo Life Sciences, BML-N156-0005, NY, USA), OR1D2 antibody (antibody epitope: 201-250 amino acids, Sigma-Aldrich, SAB4502050, St. Louis, MO, USA), rabbit polyclonal antibody against G_olf_ subunit (Santa Cruz Biotechnology, Sc-385, CA, USA), mouse monoclonal antibody against Gα_s_ (antibody epitope: 11-21 amino acids of human origin, DQRNEEKAQRE, Santa Cruz Biotechnology, Sc-135914, CA, USA), human tubal fluid (HTF) medium and human serum albumin (HSA) (Merck Millipore Corporation, Billerica, MA, USA), Fluo-4 AM and Pluronic F-127 (Molecular Probes, Eugen, OR, USA), anti-Actin and anti-GAPDH antibodies (Proteintech Group, Wuhan, China), the HRP-conjugated goat anti-rabbit IgG secondary antibody (Thermo Fisher Scientific, Waltham, MA, USA). All the other reagents were obtained from Sigma-Aldrich (St. Louis, MO, USA) unless otherwise stated.

### Sperm preparation

Human sperm were freshly obtained from healthy young donors who had reproductive history during the preceding two years. As previous described (63), liquefied human sperm were purified by direct swim-up in high-saline (HS) solution containing (in mM): 135 mM NaCl, 5 mM KCl, 1 mM MgSO_4_, 2 mM CaCl_2_, 20 mM HEPES, 5 mM glucose, 10 mM lactic acid and 1 mM Na-pyruvate, adjusted to pH 7.2 with NaOH. After being purified, semen samples were used in [Ca^2+^]_i_ measurements, sperm patch clamping and intracellular cAMP measurement. For sperm chemotaxis assay, human sperm were initially capacitated in HTF supplemented with 4 mg/ml HSA for 3 h.

### Ethical approval

The donors participating this study signed the informed consent form. The collection of semen and experiments in this study were approved by the Institutional Ethics Committee on human subjects of Jiangxi Maternal and Child Health Hospital.

### Sperm chemotaxis bioassay

Sperm chemotaxis was assessed as described (8) with some modifications. Briefly, 5 μL 1 μM bourgeonal (in HTF) and 5 μL HTF were introduced successively into 7.5-cm flattened capillary tubes (1.0-mm inner depth; Elite Medical Co., Ltd., Nanjing, China) through the same end, while the other end was sealed with plasticine. Bourgeonal concentration gradient should be formed in the capillary tubes according to the principle of fluid dynamics. In control groups, capillaries were fulfilled with HTF. To assess effects of different chemicals on human chemotaxis induced by bourgeonal, both bourgeonal and chemicals were premixed in HTF. Next, the open ends of capillaries were vertically inserted into 100 μL capcitated samples (1-5 × 10^8^ cells per ml) and incubated (37 °C, 5% CO_2_) for 1 h. Subsequently, the tubes were removed, wiped, and imaged with a Leica DM2500 Upright Microscope. Three fields at 1 and 2 cm from the base of the tube were counted, and cell densities were calculated by average cells/fields. The cell densities were employed to assess sperm chemotaxis movement and they were normalized to values from parallel, untreated controls (HTF).

### Single-sperm calcium imaging

After purification, samples were resuspended in HS and the sperm concentration was adjusted to 1-5 ×10^7^ cells per ml, then samples should be loaded by Fluo-4 AM (Molecular Probes, USA) and Pluronic F-127 (Molecular Probes, USA) in incubator (37°, 5% CO_2_) for 30 min. The fluctuation of sperm [Ca^2+^]_i_ was examined as previously described (64). Before the addition of various chemicals, sperm [Ca^2+^]_i_ was monitored in HS bath solution for 2-3 minutes until becoming stable. In parallel controls, the influence of 80 μM bourgeonal on sperm [Ca^2+^]_i_ was detected. To determine the effects of various chemicals on bourgeonal-induced sperm [Ca^2+^]_i_, sperm were pretreated with chemicals for 3-5 minutes, then bourgeonal were added to test the change in sperm [Ca^2+^]_i_. OR1D2 antibody was denatured in boiling water for 30 minutes to examine the inhibition specificity of antibodies on bourgeonal-evoked [Ca^2+^]_i_. All data were recorded and analyzed with commercial software (MetaFluorv7, Molecular Devices, Sunnyvale, CA, USA). The change of sperm [Ca^2+^]_i_ was calculated by ΔF/F_0_ (%) indicating the percent (%) of fluorescent changes (ΔF) normalized to the mean basal fluorescence before the application of any chemicals (F_0_). ΔF/F_0_ (%) = (F-F_0_)/F_0_ × 100%, F indicates the fluorescent intensity at each recorded time point.

### Western blot

After the completion of the previous steps, sperm proteins were extracted according to reference (65). The protein concentrations were determined by the bicinchoninic acid (BCA) method (Thermo Scientific, USA). The immunoblotting assays were conducted as described (63) and the dilutions of primary and secondary antibodies were as follows: 1:500 for anti-OR1D2 antibody; 1:10000 for secondary antibodies, 1:20000 for anti-GAPDH antibody. After incubation, immunoreactivity was detected using chemiluminescence detection kit reagents (Thermo Scientific, USA) and a Chimidoc™ Station (Bio-Rad).

### Human sperm patch-clamp recording

Whole-cell currents were recorded by patch-clamping the sperm cytoplasmic droplet as reported previously (63). For recording of the human CatSper current, glass pipettes (15-25 MΩ) were filled with CatSper pipette solution, which containing 130 mM caesium methane-sulphonate, 20 mM HEPES, 5 mM EGTA, 5 mM CsCl, adjusted to pH 7.2 with CsOH. After gigomal seal was formed between pipette and sperm droplet in HS bath solution, the transition into whole-cell mode was made by application of short (1 ms) voltage pulses (400-650 mV) combined with light suction. The baseline CatSper currents were recorded in HS solution. Subsequently, monovalent currents of CatSper could be detected when perfusing sperm with divalent-free solution (NaDVF), which containing 150 mM NaCl, 20 mM HEPES, 5 mM EDTA, adjusted to pH 7.2 with NaOH. To test the effects of various chemicals on bourgeonal-induced CatSper currents, bourgeonal and other chemicals was premixed in NaDVF. The effect of G proteins on bourgeonal-induced CatSper currents was examined by intracellular application of GTP, GDP-β-S, anti-G_olf_ and anti-Gα_s_ antibodies. All patch-clamp data were analyzed with Clampfit version 10.4 software (Axon, Gilze, Netherlands).

### Measurement of the cAMP level in human sperm

After purification was completed, human sperm were resuspended in HS solution and the sperm concentration was adjusted to 1-5 × 10^8^ cells per ml by using computer-assisted semen analysis (CASA) system (WLJY-9000, WeiLi. Co., Ltd., Beijing, China). The cell suspension would be divided equally into different tubes (100 μL per tube). In control, intracellular cAMP concentration of sperm in HS (with 0.1% DMSO) were detected. To assess the effects of various reagents on sperm cAMP, bourgeonal and other chemicals were incubated with sperm in incubator (37ଌ, 5% CO_2_) for a certain period. 0.1 mM HCl was added into the tubes to terminate reactions and intracellular cAMP was liberated in ice bath by ultrasonic treatments (400W, 4s ultrasonic time, 8s time interval, 3 minutes). Afterwards, these tubes should be agitated in ice bath for 2h, then the cell lysate should be diluted at least two times by ELISA buffer (cAMP assay kit, 581001, Cayman Chemical). Subsequently, the instructions of the kit were followed. Finally, the absorbance of each plate was read at the wavelength of 415 nm and absorbance data were analyzed by employing the computer spreadsheet, which could be download from www.caymanchem.com/analysis/elisa freely.

### Statistical analysis

Data were expressed as the mean ± SEM. Shapiro-wilk test was utilized to test for normality of data distribution. Worthy of note, the differences between the data groups, which did not coincide with normal distribution, were analyzed by Mann-Whitney test, a kind of non-parametric test. In contrast, parametric tests (such as paired t test, etc) are more effective than non-parametric tests in analyzing the differences between the normal distribution data groups while adding directional errors when analyzing non-normal data. Newman-Keuls Multiple comparison test is a range test that ranks group means and computes a range value, and it was utilized to analyze statistical differences between control and experimental groups in Figure 5A and B; Figure S6B; Figure S13C. Statistical differences were determined by the statistical software GraphPad Prism (version 5.01, GraphPad Software, San Diego, CA, USA). Statistical significance is expressed as follows: not significant (ns), *\*P* < 0.05, *\*\*P* < 0.01, *\*\*\*P* < 0.001.

Additional experimental procedures are in *Appendix supplementary Materials and Methods*.

## Acknowledgement

The authors declare no conflict of interest. We appreciate Tao Wang’s contribution when initiating this work and his considerable thoughts regarding the manuscript. We thank Dr. Christopher J Lingle and Dr. Jin Zhang for helpful suggestions with regard to this article. This work was supported by research grants from National Basic Research Program of China (973 Program, No. 2015CB943003) and National Natural Science Foundation of China (No. 31671204 & No. 31230034) to Xu-Hui Zeng.

## Author contributions

XHZ conceived the project. YMC, TL, ZP, designed and performed experiments. HYC acquired and processed all semen samples. XHZ and YMC wrote and revised the manuscript.

